# The structure of the influenza A virus genome

**DOI:** 10.1101/236620

**Authors:** Bernadeta Dadonaite, Egle Barilaite, Ervin Fodor, Alain Laederach, David L. V. Bauer

**Affiliations:** Sir William Dunn School of Pathology, University of Oxford, South Parks Road, Oxford OX1 3RE, UK; Department of Biology, University of North Carolina, Chapel Hill, North Carolina, USA

## Abstract

Influenza A viruses (IAVs) are segmented single-stranded negative sense RNA viruses that constitute a major threat to human health. The IAV genome consists of eight RNA segments contained in separate viral ribonucleoprotein complexes (vRNPs) that are packaged together into a single virus particle^1,2^. While IAVs are generally considered to have an unstructured single-stranded genome, it has also been suggested that secondary RNA structures are required for selective packaging of the eight vRNPs into each virus particle^3,4^. Here, we employ high-throughput sequencing approaches to map both the intra and intersegment RNA interactions inside influenza virions. Our data demonstrate that a redundant network of RNA-RNA interactions is required for vRNP packaging and virus growth. Furthermore, the data demonstrate that IAVs have a much more structured genome than previously thought and the redundancy of RNA interactions between the different vRNPs explains how IAVs maintain the potential for reassortment between different strains, while also retaining packaging selectivity. Our study establishes a framework towards further work into IAV RNA structure and vRNP packaging, which will lead to better models for predicting the emergence of new pandemic influenza strains and will facilitate the development of antivirals specifically targeting genome assembly.

---

Influenza A viruses cause seasonal epidemics as well as occasional pandemics. The segmented nature of the IAV genome allows reassortment of viral genome segments between established human influenza viruses and influenza viruses harboured in the animal reservoir^5^. This can lead to emergence of novel influenza strains, against which there is little pre-existing immunity in the human population^6^. However, due to a lack of understanding of the molecular mechanisms governing the packaging of the eight genome segments into a single virion, it remains unclear which influenza virus strains have the potential to form reassortants. In virions, as well as in infected cells, the RNA genome segments are assembled into viral ribonucleoprotein (vRNP) complexes in which the termini of the viral RNA (vRNA) associate with the viral RNA-dependent RNA polymerase while the rest of the vRNA is bound by oligomeric nucleoprotein (NP)^2,7^. Although cryo-EM studies revealed the overall architecture and organisation of vRNPs, the resolution of currently available structures is not sufficiently high to provide information about the conformation of the vRNA^8,9^. It is thought that, through specific RNA-RNA interactions, exposed regions of vRNA in the vRNP mediate segment-specific interactions during virion assembly, ensuring that the correct set of eight vRNPs is selected^3,4^. However, the identities of these interacting regions as well as the overall structure of vRNA in vRNPs are currently unknown.

To better understand the vRNA structure in the context of vRNPs, we employed SHAPE-MaP (Selective 2’-Hydroxyl Acylation Analysed by Primer Extension and Mutational Profiling), which probes the conformational flexibility of each vRNA nucleotide both *ex virio* and *in virio* (**Extended Data Fig. 1**)^10,11^. For the *ex virio* experiments, the eight vRNA segments were individually transcribed from plasmid DNA using T7 RNA polymerase (ivtRNA) or “naked” vRNA was purified from deproteinated influenza A/WSN/1933 (H1N1) (WSN) particles (nkvRNA). For the *in virio* experiments, vRNA was probed in the context of vRNPs directly inside purified virions (**Extended Data Fig. 1a**).

Three biological replicates of the *in virio* SHAPE-MaP experiments resulted in highly reproducible SHAPE reactivity profiles suggesting that there is an underlying structure of vRNA in the context of vRNP (**Extended Data Fig. 2a**). We found that the eight different vRNPs *in virio* have unique vRNA conformations, as demonstrated by the different SHAPE reactivity profiles (**Fig. 1a**). Regions of extensive low SHAPE reactivities *in virio* (where the median-SHAPE value is below zero, **Fig. 1a**) indicate that the vRNA in the context of vRNPs is capable of accommodating secondary RNA structures with extensive base-pairing. However, comparison of *in virio* and *ex virio* SHAPE profiles shows a significant shift in the distribution of SHAPE reactivities in virio, consistent with an overall more open (less structured) RNA conformation (**Fig. 1b**). In addition, the vRNA forms fewer high-probability secondary RNA structures *in virio*, suggesting that the binding of NP remodels and partially melts secondary structures in vRNPs (Fig. 1c), in agreement with early studies using enzymatic and chemical probing of naked and NP-bound short RNAs^12^. Regions of high correlation between *ex virio* and *in virio* SHAPE profiles reveal regions of vRNA which are capable of local secondary structure formation even when NP is bound. Our data also recapitulate the RNA structures that have been identified previously using computational methods^13,14^ (**Fig. 1d-e; Extended Data Fig. 2b; Extended Data Fig. 3-4**).

**Figure 1.**
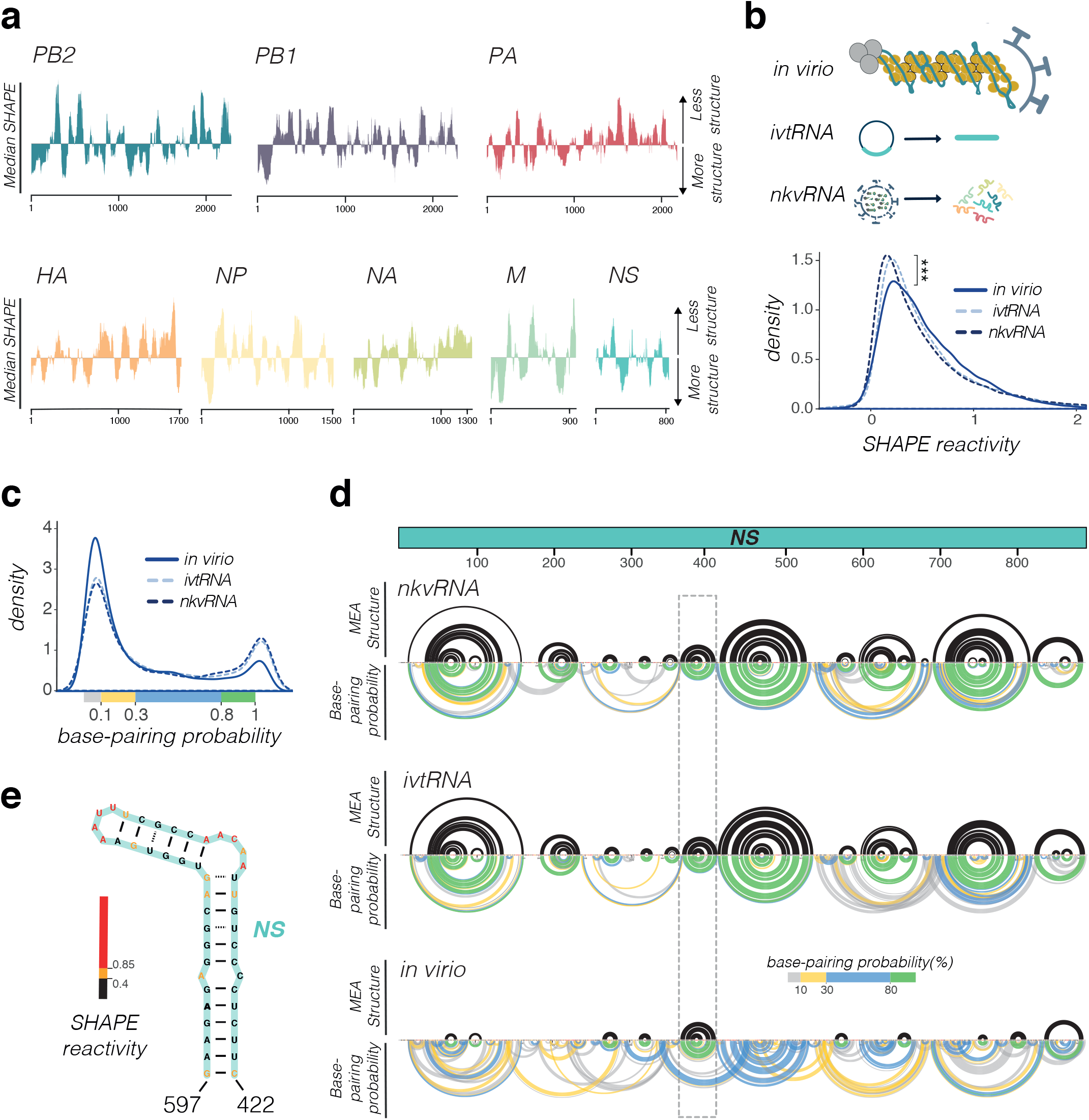
SHAPE-MaP analysis of the influenza A virus genome structure. **a,** Median SHAPE reactivities of different vRNA segments. Medians were calculated over 50 nucleotide windows and plotted relative to the global median. **b,** vRNA SHAPE reactivity distributions in different samples; ***P<0.0001, Wilcox Rank-Sum Test. **c,** Base-pairing probability distributions in different vRNA samples; **d,** Secondary RNA structure of the NS segment. Black arcs indicate the maximum expected accuracy RNA structure; only the interactions associated with greater than 80% base-pairing probabilities are shown. Coloured arcs indicate the base-pairing probabilities. Dashed rectangle highlights the position of the hairpin shown in **e.** All sequence positions are annotated as 5’-3’ in vRNA sense. nkvRNA, naked viral RNA; ivtRNA, in vitro transcribed RNA; MEA, maximum expected accuracy.

The finding that secondary RNA structures are accommodated in vRNPs suggests that NP is not distributed on the vRNA uniformly, in agreement with a recent study of NP association with vRNA^15^. This raises the possibility that parts of the vRNA could be exposed and accessible to form intermolecular RNA-RNA interactions. Indeed, it has been previously proposed that the specific packaging of the eight different vRNPs into the budding virus is mediated by selective RNA-RNA interactions forming between the vRNPs^16-21^. We therefore proceeded to analyse the intermolecular RNA interactions occurring *in virio* using SPLASH (Sequencing of Psoralen Crosslinked, Ligated, and Selected Hybrids)^22^. SPLASH uses a reversible intercalating reagent, psoralen, which crosslinks base-paired RNAs and allows mapping and identification of crosslinked regions using high-throughput sequencing. We preformed two biological replicates of SPLASH analysis using purified virions and focussed our analysis on the most prevalent RNA interactions found in both experiments (**Extended Data Fig. 5**). 84% of interactions were observed in both replicates.

Our analysis shows that the distribution of intermolecular interaction sites varies between the eight different vRNA segments and interactions sites are not restricted to certain regions, e.g. vRNA termini (**Fig. 2a**). Most segments can interact with multiple other segments and in some cases the same region can mediate interactions with multiple segments. While it is unlikely that the same region would interact with multiple other segments in the same virion, this finding suggests that certain loci in the vRNA are more likely to be involved in intermolecular RNA base-pairing than others. These data also suggest that there is a level of redundancy in the intermolecular interactions, allowing multiple RNA conformations to be packaged *in virio*. Notably, when we used an intermolecular RNA interaction algorithm to compute the intermolecular free energy of the RNA-RNA interaction sites identified by SPLASH, we saw a significant enrichment in low energy (highly favourable interactions) compared to random permutations of the same interaction dataset (**Fig. 2b-c**). Furthermore, SHAPE-informed intramolecular RNA interaction prediction shows that the identified interactions are compatible with the *in virio* SHAPE reactivity profiles we have determined previously and indicates that the low free energy of the interactions is maintained (**Extended Data Fig. 6**).

**Figure 2.**
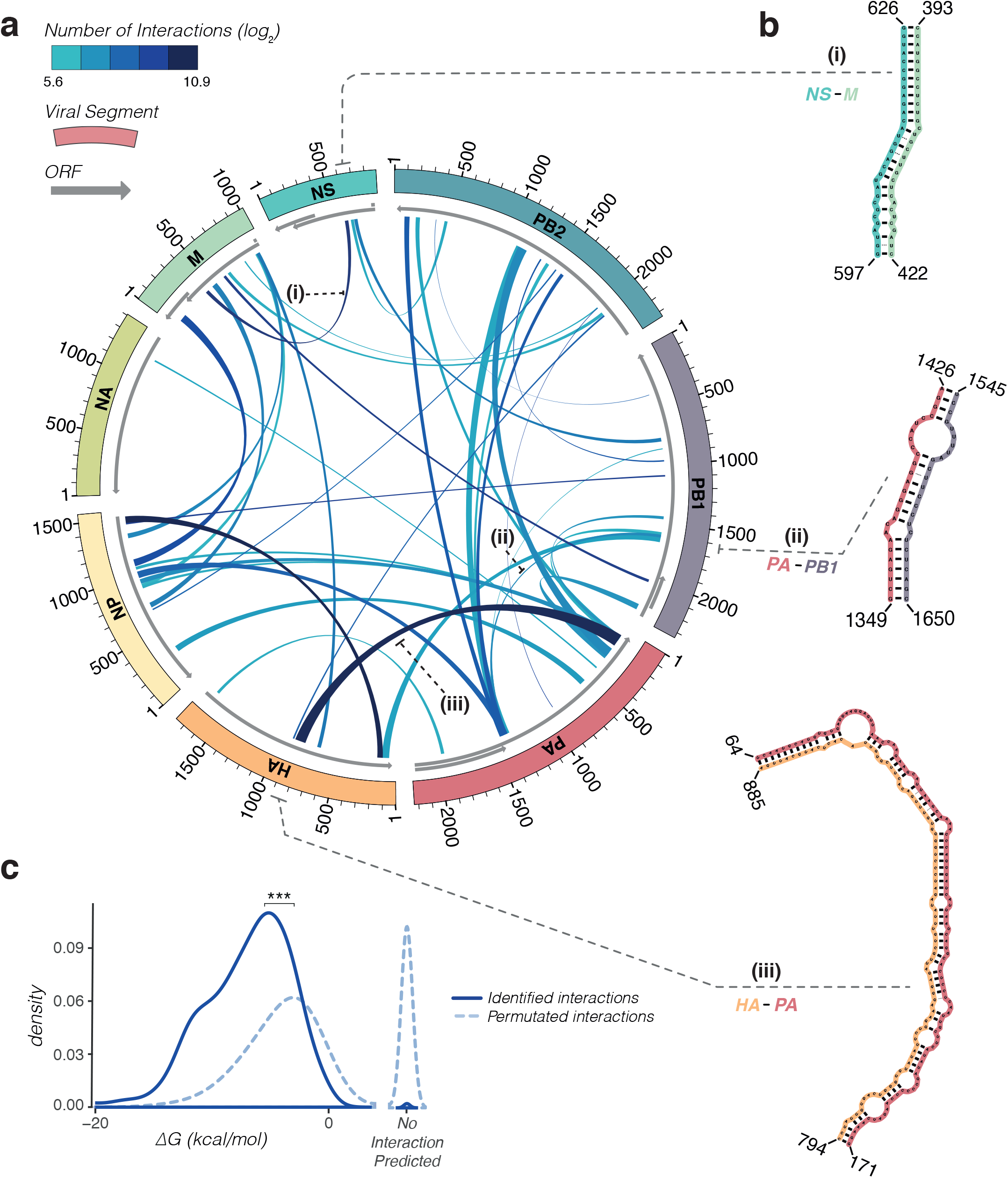
Intermolecular RNA interactions in the IAV genome. **a,** The most common intermolecular RNA interactions identified using SPLASH. The links indicate the regions involved in inter-segment base-pairing and are coloured by interaction depth on a log2 scale. The top 8% of interactions are shown, which account for 82% of total chimeric reads identified. **b,** Examples of the identified intermolecular interaction structures. **c,** Probability distributions of the ΔG energy distributions associated with the interactions identified by SPLASH versus a permutated interaction dataset (n=473). ***P<0.00001, Wilcox Rank-Sum Test.

Previous studies suggested that the eight vRNA segments are assembled in a hierarchal manner, with some segments being more critical than others^1^. Specifically, it was found that the NA and NS segments are the most easily exchangeable between viral strains, suggesting that they make the least contribution to the hierarchal assembly of the eight vRNPs^23-25^. In agreement, we observe that NA and NS segments form the fewest interactions with other segments. Surprisingly, we also identify very few interactions between the NA and HA segments (0.13% of the mapped reads), suggesting that influenza viruses maintain the greatest possible antigenic diversity by limiting the interactions between these two segments during genome packaging.

The sequence of the IAV genome undergoes changes as a result of the antigenic drift and shift^5^. Given that the intermolecular RNA interactions we have identified may be important for reassortment between the different IAV strains, we questioned whether the same interactions could occur in other IAV strains as well. We selected a set of IAV strains representing the pandemic strains of the last century (A/Brevig Mission/1/1918 (H1N1), A/Singapore/1/57 (H2N2), A/Hong Kong/1/68 (H3N2), and A/England/195/2009(H1N1)), and analyzed their potential to form intermolecular RNA-RNA interactions in the same regions we identified in WSN. Though not all of the interactions can form in the different strains, the regions corresponding to those we identified in WSN are more likely to be involved in the intermolecular base-pairing than permutated datasets (**Fig. 3a**), and a number of extensive interactions are maintained in the different viral strains (**Fig. 3b**).

**Figure 3.**
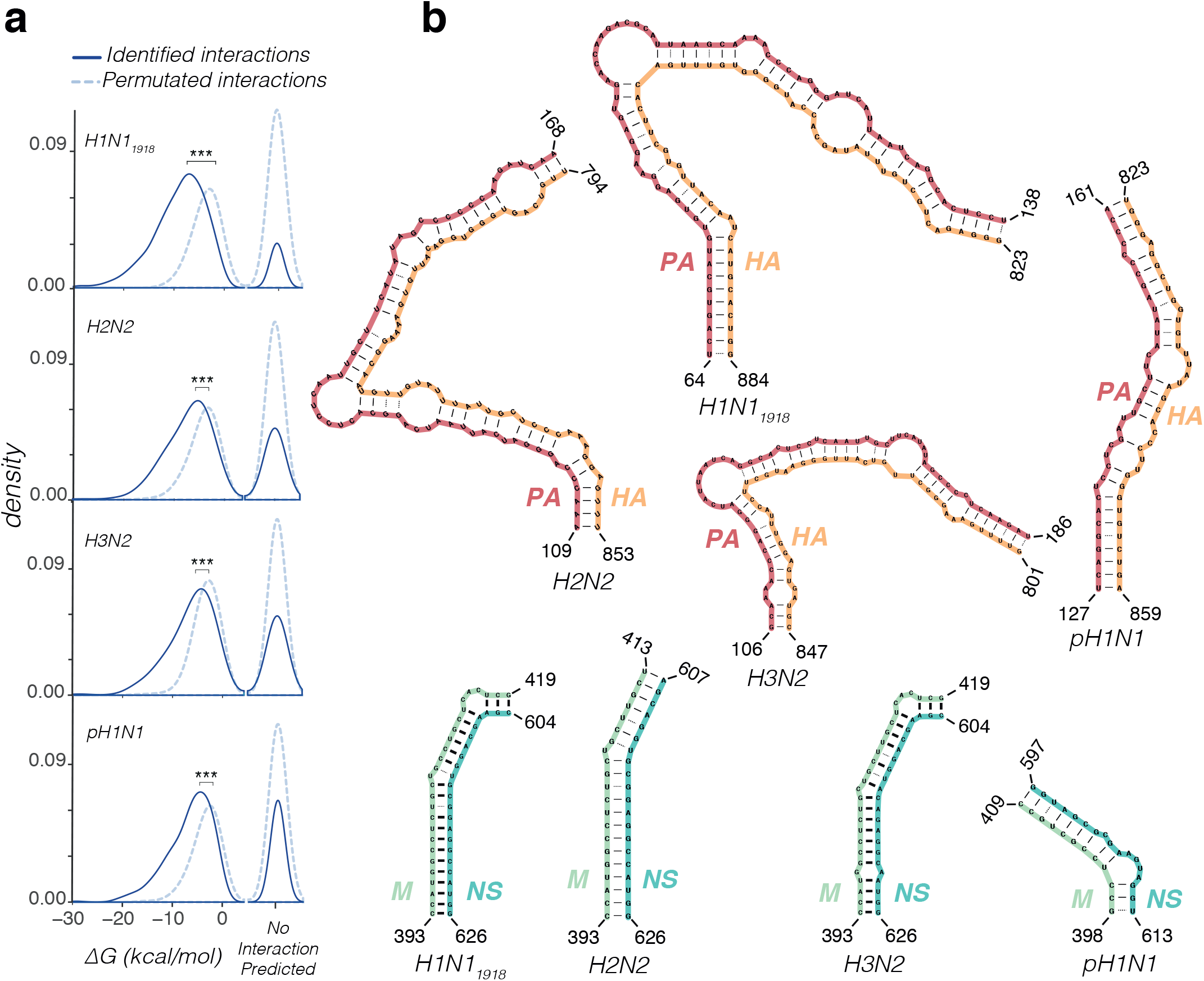
Intermolecular RNA interactions are conserved in different IAV strains. **a,** Distribution of ΔG energies associated with the intermolecular RNA interactions in different IAV strains; ***P<0.0001, Wilcox Rank-Sum Test. **b,** Examples of conserved intermolecular interaction structures. H1N11918, A/Brevig Mission/1/1918 (H1N1); H2N2, A/Singapore/1/57 (H2N2); H3N2, A/Hong Kong/1/68 (H3N2); pH1N1, A/England/195/2009 (H1N1).

To address the biological role of the identified *in virio* RNA-RNA interactions we used synonymous mutagenesis to disrupt RNA interactions while preserving the encoded amino acid sequence. We find that mutant viruses with decreased strength of intermolecular RNA-RNA interactions have significant differences in the ratios of the different segments packaged into the virions compared to both the wild type virus and to a control virus with mutations which do not interfere with identified intermolecular RNA-RNA interactions (**Fig. 4a**). In addition, weakening of intermolecular RNA-RNA interactions leads to the production of defective viral particles and changes in the kinetics of virus growth (**Fig. 4b-c**).

**Figure 4.**
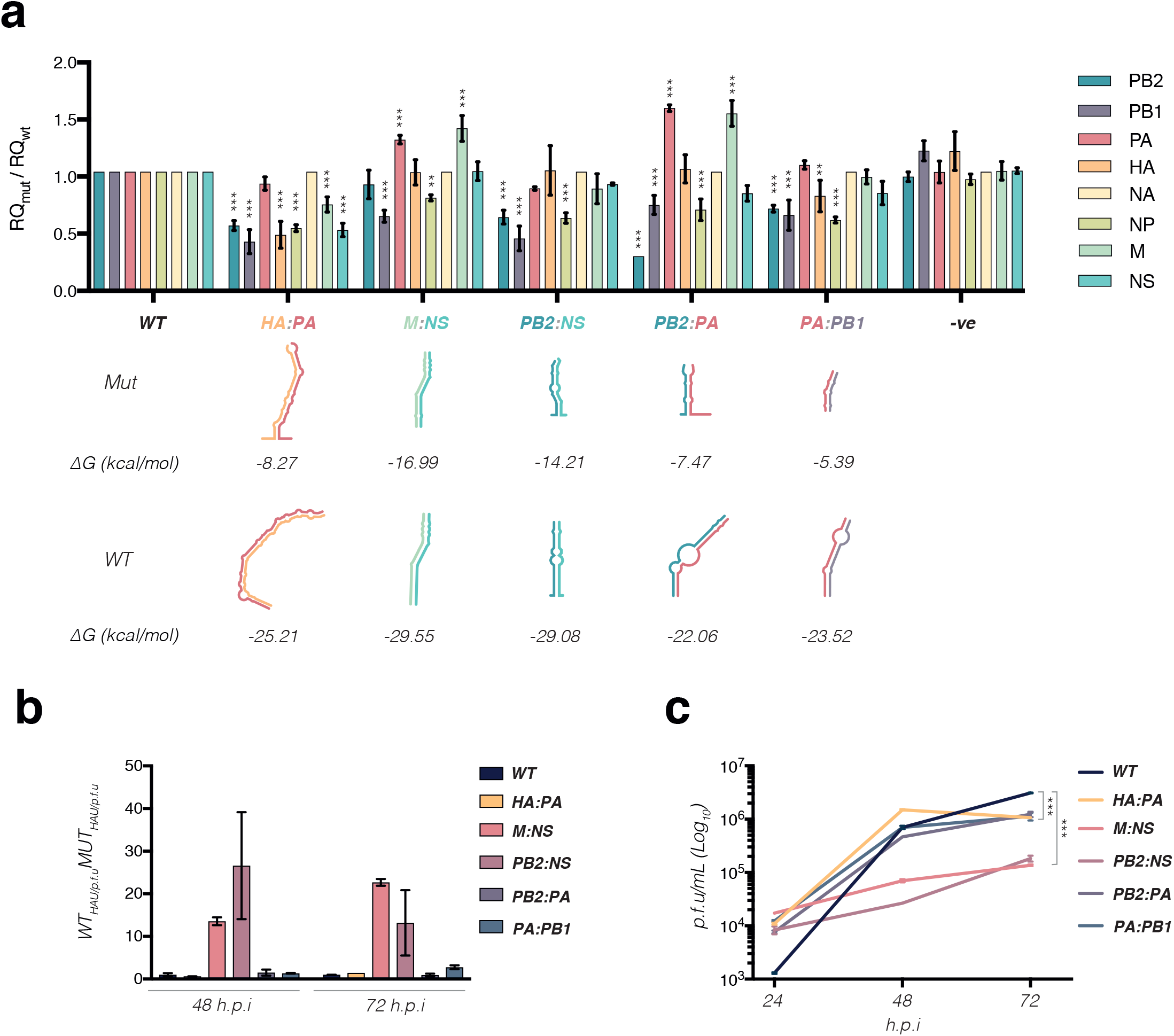
Intermolecular RNA interactions are important for the packaging of vRNPs. **a,** RT-qPCR analysis of the vRNA inside viruses with disrupted intermolecular interaction sites. For each virus an RQ value relative to the NA segment was calculated and the changes compared to the wild-type virus are shown. Error bars indicate standard deviation; data from three technical replicates is shown. **b,** Quantification of HAU to p.f.u. ratios in mutant viruses relative to the wild type. Quantification was done on the 48 and 72 h.p.i samples from the virus growth analysis shown in **c. c,** Growth kinetics of mutant viruses. Error bars indicate standard deviation; data from two biological replicates are shown. **P<0.001, ***P<0.0001, ANOVA with Dunnett’s Multiple Comparison Test. RQ, relative quantification; p.f.u., particle forming units; h.p.i., hours post infection. Mutant viruses are labelled by indicating disrupted segment interactions: HA:PA, HA:794-885 PA:64-171; M:NS, M:393-422 NS:597-626; PB2:NS, PB2:437-462 NS:649-674; PB2:PA, PB2:89-132 PA:1406-1447; PA:PB1, PA:1375-1400 PB1:1603-1627.

Overall, we present the first global map of the IAV genome structure *in virio*. We show that the IAV genome contains both intramolecular and intermolecular RNA structures. Importantly, our study shows that *in virio*, IAV maintains an extensive set of RNA-mediated interactions between vRNPs, which is important for the packaging of the viral genome. Maintaining a redundant inter-vRNP interaction network to facilitate the selective packaging of the different genomic segments (vs. a limited set of interactions), could be a strategy to balance the need for selective packaging with the ability to allow reassortment to occur. Our analysis suggests that some, though not all, of the identified intermolecular interactions are present in evolutionarily distant IAVs. A redundant inter-vRNP interaction network could allow multiple pathways towards assembling the full set of eight vRNPs; having multiple pathways is potentially important for the emergence of novel influenza virus strains through reassortment of genome segments of two evolutionarily distant viruses.

This evolutionary flexibility of genome assembly is also apparent in the location of specific intermolecular RNA-RNA interaction regions. A number of these overlap with regions of the genome that are evolutionarily less constrained with respect to their protein-coding capacity. For example, highly prevalent interactions of the NA and NS segments fall into regions encoding the NA stalk and the linker between the N-terminal RNA-binding domain and the C-terminal effector domain of NS1, respectively (**Extended Data Fig. 7**). Furthermore, a prominent interaction hotspot in the PA segment, involved in interactions with multiple other segments, lies immediately downstream of the overlapping PA-X open reading frame (ORF) in a region that encodes the linker between the N-terminal endonuclease and C-terminal domain of PA. These observations suggest that the positioning of some of the intersegment RNA interaction sites may be constrained by the balance between the constantly drifting RNA sequences and maintenance of the encoded amino acids.

In virions, the vRNPs are organized in a “7+1” pattern with seven segments of different lengths surrounding a central segment^17,26-28^. This model has led to the suggestion that the central segment may act as a ‘master segment’ mediating the selection of the other segments. We note that our analysis shows that the PA segment is capable of forming multiple strong intermolecular interactions, with the 5’ end and the 1400-1500 nt region involved in multiple redundant interactions with many other segments.

We anticipate that further studies of IAV genome structure and RNA-RNA interaction networks will lead to an improved ability to predict the potential for reassortment between different IAV strains and consequently will facilitate the prediction of the emergence of new pandemic influenza strains. Furthermore, such studies may guide the design of new antivirals targeting the assembly of the eight vRNPs and blocking virus packaging.

## Acknowledgements

We thank J. Kenyon for helpful discussions and sharing protocols and J. Robertson for making 1M7 reagent. This work was supported by a Wellcome Trust studentship [105399/Z/14/Z] (to B.D.), a Medical Research Council programme grant [MR/K000241/1] (to E.F.), and an EPA Cephalosporin Junior Research Fellowship (to D.L.V.B.). This work was also supported by the U.S. National Institutes of Health [grant numbers HL111527, GM101237 and HG008133].

## Contributions

B.D. Designed and preformed the experiments, analysed the data and wrote the paper; E.B. made mutated viruses. E.F. supervised virus work and edited the manuscript; A.L. supervised SHAPE data analysis and RNA modelling. D.V.L.B. designed SPLASH experiments, analysed data, supervised sequencing work and edited the manuscript.

## Author Information

Sequencing data have been deposited to Sequence Read Archive (accession SRP127020 & SRP126994) and the processed SHAPE reactivities and SPLASH data are available in SNRNASM format as a **supplemental table 1**. The authors declare no competing financial interests. Correspondence and requests for materials should be addressed to E.F. and D.L.V.B.

## Methods

### Cell culture, virus growth and purification

Madin-Darby Bovine Kidney (MDBK) epithelial cells were grown in Minimum Essential Medium (MEM; Merck), supplemented with 2 mM L-glutamine and 10% fetal calf serum. Human embryonic kidney 293T (HEK 293T) cells were maintained in Dulbecco’s Modified Eagle Medium (DMEM; Merck). Viral stocks were produced by infecting MDBK cells with influenza A/WSN/33 (WSN) (H1N1) virus at an MOI of 0.001. Virus was harvested 2 days post infection. Virus stocks were purified by ultracentrifugation: firstly, the infected cell culture medium was clarified by centrifugation at 4000 rpm for 10 min at 4°C followed by centrifugation at 10,000 rpm for 15 min at 4°C. The virus was then purified by centrifugation through a 30% sucrose cushion at 25,000 rpm for 90 min at 4°C in a SW32 rotor (Beckman Coulter). The purified virus pellet was resuspended in a resuspension buffer (0.01 M Tris-HCl (pH 7.4), 0.1 M NaCl, 0.0001 M EDTA). Viruses containing synonymous mutations were produced using the 12-plasmid rescue system as described previously ^29^. The primers used to generate mutations are provided in **Extended Data Table 1**. Virus growth curves were generated by infecting a 70% confluent MDBK cell layer with viral stocks at an MOI of 0.001. Supernatant from infected cells was collected at 24, 48 and 72 h post infection. The infectious virus titres were determined by plaque assay. The significance of the differences between mutated and wild type virus growth kinetics was assessed using ANOVA with Dunnett’s Multiple Comparison Test on GraphPad Prism software. Haemagglutination assays were carried out by serially-diluting the supernatants from infected cells in phosphate-buffered saline (PBS) in a 96-well plate. An equal volume of 0.5% chicken blood was added to the serial dilutions and the plates were incubated at 4°C until hemagglutination was observed.

### Selective 2’-hydroxyl acylation analysed by primer extension and mutational profiling (SHAPE-MaP)

1-methyl-7-nitroisatoic anhydride (1M7) was custom synthesised from 4-nitroisatoic anhydride as described previously^30^. For the ivtRNA experiments each vRNA segment was synthesised from a linear DNA template using the HiScribe^^™^^ T7 High Yield RNA Synthesis Kit (NEB). The products were checked for size and purity on a 3.5% PAGE-urea gel. nkvRNA samples were prepared by purifying the WSN particles over sucrose cushion as described above. Purified viruses were treated with 250 μg/mL of Proteinase K (Roche) in PK buffer (10 mM Tris-HCl (pH 7.0), 100 mM NaCl, 1 mM EDTA, 0.5% SDS) for 40 min at 37°C. Before the modification ivtRNA and nkvRNA samples were folded at 37°C for 30 min in folding buffer (100 mM Hepes-NaOH (pH 8.0), 100 mM NaCl, 10 mM MgCl_2_. 1M7 (dissolved in anhydrous DMSO (Merck)) was added to a final concentration of 10 mM to the folded RNA and the samples were incubated for 75 s at 37°C. The *in virio* modifications were performed by adding 1M7 directly to the purified virus stocks. The ability of SHAPE reagents to penetrate viral particles was initially tested as described previously ^31^ by preforming ^32^P-labelled primer extensions on RNA extracted from SHAPE reagent-treated viral stocks using an NA segment targeting primer (5’-AATTGGTTCCAAAGGAGACG-3’). In parallel to the 1M7-treated samples, control samples were treated with DMSO. RNA extracted from purified viral particles or denatured T7 RNA polymerase transcribed RNA was used for denatured controls (DC). To prepare DC samples the RNA was mixed in DC buffer (50 mM Hepes-NaOH (pH 8.0), 4 mM EDTA) with 55 % formamide and incubated at 95°C for 1 min. 1M7 was then added to 10 μM and the samples were incubated at 95°C for an additional 1 min. N-methylisatoic anhydride (NMIA, Thermo Fisher) SHAPE reagent was also tested *in virio*. Experiments with NMIA were preformed as described above for 1M7, except the purified virions were treated with NMIA for 45 min.

Sequencing library preparation was done as described previously^11^ following the randomer workflow. In brief, after 1M7 or control treatments, RNA was cleaned up using the RNA Clean & Concentrator^™^-5 kit (Zymo Research). The RNA was reverse transcribed using Random Primer Mix (NEB) with Superscript II in MaP buffer (50 mM Tris-HCl (pH 8.0), 75 mM KCl, 6 mM MnCl_2_, 10 mM DTT and 0.5 mM dNTPs). Nextera XT DNA Library Prep Kit (Illumina) was used to prepare the DNA libraries. Final PCR amplification products were size selected using Agencourt AMPure XP beads (Beckman Coulter) and quality assessed using the Agilent DNA 1000 kit on a Bioanalyser 2100 instrument (Agilent). The libraries were sequenced (2x150bp) on a HiSeq4000 instrument (Illumina).

### Sequencing of psoralen crosslinked, ligated, and selected hybrids (SPLASH)

SPLASH samples were prepared as published previously^22,32^ with some modifications. Purified virus stocks were incubated with 200 μM of EZ-Link^™^ Psoralen-PEG3-Biotin (Thermo Fisher) and 0.01% digitonin (Merck) for 5min at 37°C. The viruses were spread on a 6-well dish, covered with a glass plate, placed on ice, and irradiated for 45 min using a UVP Ultra Violet Product™ Handheld UV Lamp (Fisher). Cross-linked virus stock was treated with Proteinase K (Merck) and the viral RNA (vRNA) was extracted using TRIzol (Invitrogen). An aliquot of extracted vRNA was used to detect biotin incorporation using chemiluminescent nucleic acid detection module kit (Thermo Fisher) on Hybond-N+ nylon membrane (GE Healthcare Life Science). The rest of the extracted vRNA was fragmented using NEBNext^®^ Magnesium RNA Fragmentation Module (NEB), and size selected for fragments below 200nt using RNA Clean & Concentrator^™^-5 (Zymo Research). The samples were enriched for biotinylated vRNA using Dynabeads MyOne Streptavidin C1 beads (Life Technologies) and on-bead proximity ligation and psoralen crosslink reversal were carried out as published previously. Sequencing libraries were prepared using adaptor ligation as described^22,32^ for the first SPLASH experiment, and using the commercial SMARTer smRNA-Seq Kit (Clontech) for the second SPLASH experiment. Final size selection was done by running the PCR-amplified sequencing libraries on a 6% PAGE gel (Thermo Thermo Fisher Fisher) in TBE and selecting for 200-300bp DNA. Libraries were sequenced either 1x or 2x150bp on a NextSeq 500 instrument (Illumina).

### Processing of SHAPE-MaP sequencing reads

The sequencing reads were trimmed to remove adaptors using Skewer^33^. The SHAPE reactivity profiles were generated using the published ShapeMapper pipeline^11^, which aligns the reads to the reference genome using Bowtie 2 and calculates mutation rates at each nucleotide position. The mutation rates are then converted to the SHAPE reactivity values defined as:

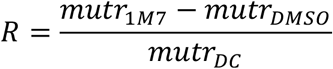

where *mutr_1M7_* is the nucleotide mutation rate in 1M7 treated sample, *mutr_1M7_* is the mutation rate in the DMSO treated sample and *mutr_DC_* is the mutation rate in the denatured control. All SHAPE reactivities are normalised to an approximate 0-2 scale by dividing the SHAPE reactivity values by the average reactivity of the 10% most highly reactive nucleotides after excluding outliers (defined as nucleotides with reactivity values that are greater than 1.5x the interquartile range).

### Processing of SPLASH sequencing reads

The sequencing reads were first deduplicated using clumpify.sh (BBMap package; https://sourceforge.net/projects/bbmap/) and adaptors were trimmed using Cutadapt^34^. STAR^35^ was used to align the reads to the WSN viral reference genome. Only the chimeric reads in which at least 30 nucleotides aligned to the reference segments were used in further processing (STAR parameter –chimSegmentMin 30). CIGAR strings in each read alignment were processed to find the read start and end coordinates. The reads aligning to the same partner segments and overlapping positions in the first and second SPLASH experiment were combined. Overlapping reads between the same partner segments in the final read set were merged and expanded to cover the total read window. The start and end coordinates for all interaction sites were defined as the 5’ and 3’ terminal positions of the expanded read site. The set of interactions was visualised using Circos^36^. Final chimeric read set is provided in **Supplementary table 2**.

### RNA structure predictions

The IntaRNA (v2.0.4) algorithm^37^ with the minimum seed requirement of 4 bp was used to predict the ability of RNA-RNA interactions to occur in the regions identified during the SPLASH analysis. Permutated data sets were generated by randomly shuffling the specific interaction partners identified by SPLASH and assessing the interaction ΔG energies using IntaRNA. The significance of the difference between the probability distributions of the ΔG energies associated with the SPLASH-identified intermolecular RNA interactions versus the permutated datasets was calculated using Wilcox Rank-Sum Test in R software. The IntaRNA structure predictions were then used to trim the interaction regions to the nucleotides involved in the base-pairing. For SHAPE-informed RNA-RNA interaction predictions RNAcofold from ViennaRNA package (v2.4.1) was used^38^. For intramolecular RNA structure predictions RNAStructure package (v6.0) was employed ^39^ using the *Fold* and *Partition* commands to predict secondary RNA structures and partition functions for each segment, respectively. SHAPE reactivities were included as pseudoenergy restraints. A 50 nt sliding median window correlation analysis between the *ex virio* and *in virio* SHAPE reactivity profiles was used to determine the extent of SHAPE correlation between the T7-transcribed RNA and vRNP-associated RNA. We found that no correlation existed >150 nt, and therefore set the maximum pairing distance constraint for structure and partition function predictions to 150 nt. For the intramolecular structure predictions we set the nucleotides within the promoter region to be single stranded.

### RT-qPCR

vRNA was extracted from rescued virus stocks using the Direct-zol^™^ RNA MiniPrep kit (Zymo Research), including a DNase treatment step. vRNA was reverse transcribed using SuperScript^™^ III (Thermo Fisher), per the manufacturer’s instructions, using an equimolar ratio of the universal vRNA primers (**Extended Data Table 2**) and primer extension at 37°C for 1 h. qPCR was performed using the Brilliant III Ultra-Fast Probe High ROX QPCR Master Mix (Agilent) with segment-specific primers and probes (**Extended Data Table 2**).

---

**Extended Data Figure 1.**
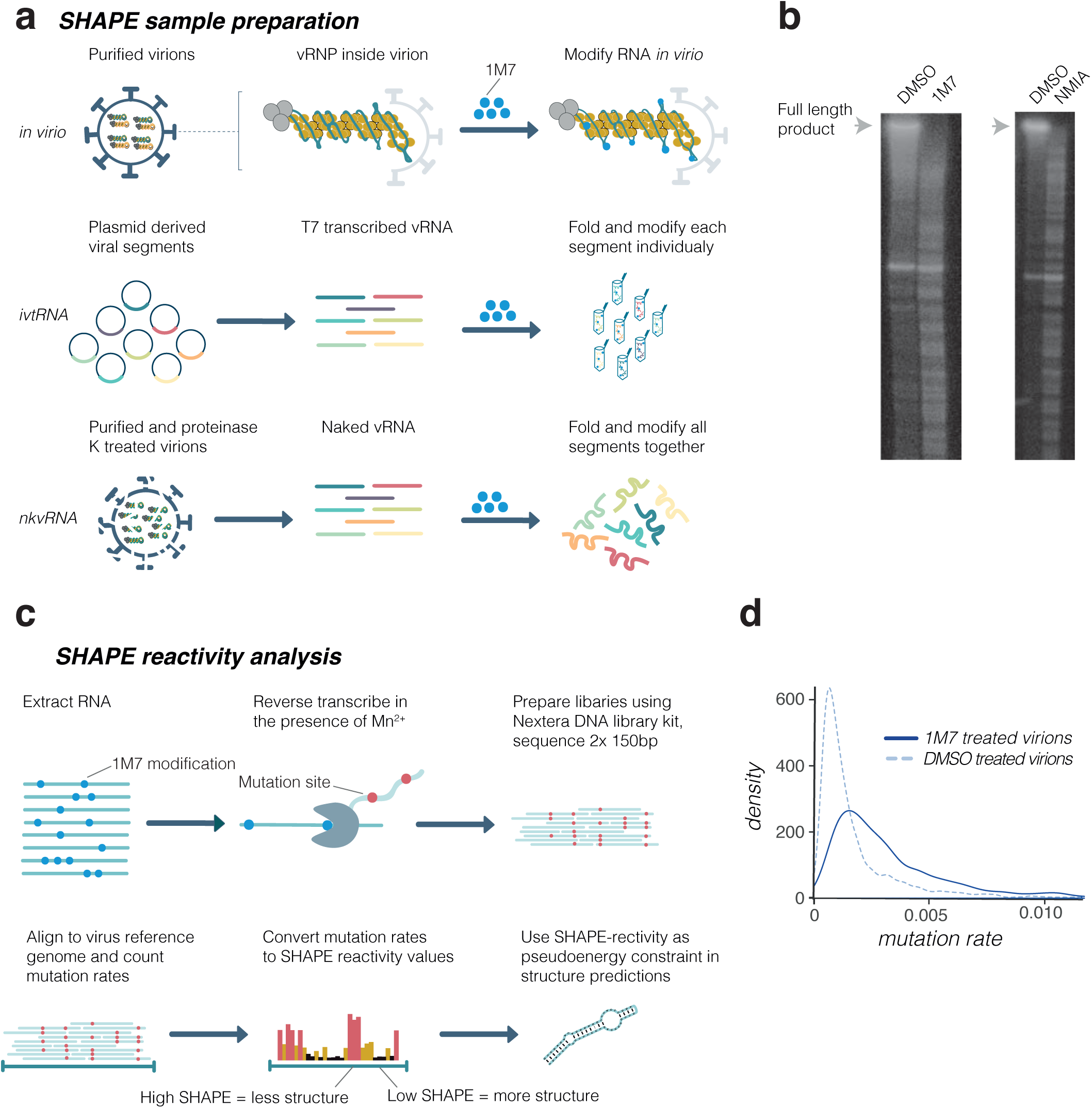
SHAPE-MaP analysis of the IAV genome. **a,** Schematic showing the vRNA samples used for SHAPE-MaP analysis. **b,** Reverse transcription reaction using vRNA that was extracted from SHAPE reagent (1M7 or NMIA) or DMSO treated viral samples. ^32^P-labelled primer targeting NA segment vRNA was used. Bands indicate the stalling of reverse transcriptase at the sites of SHAPE modifications. **c,** Schematic of SHAPE-MaP library preparation and sequencing data analysis. **d,** Mutation rates in DMSO versus 1M7 reagent treated samples. 1M7, 1-methyl-7-nitroisatoic anhydride; NMIA, N-methylisatoic anhydride; DMSO, dimethyl sulfoxide.

**Extended Data Figure 2.**
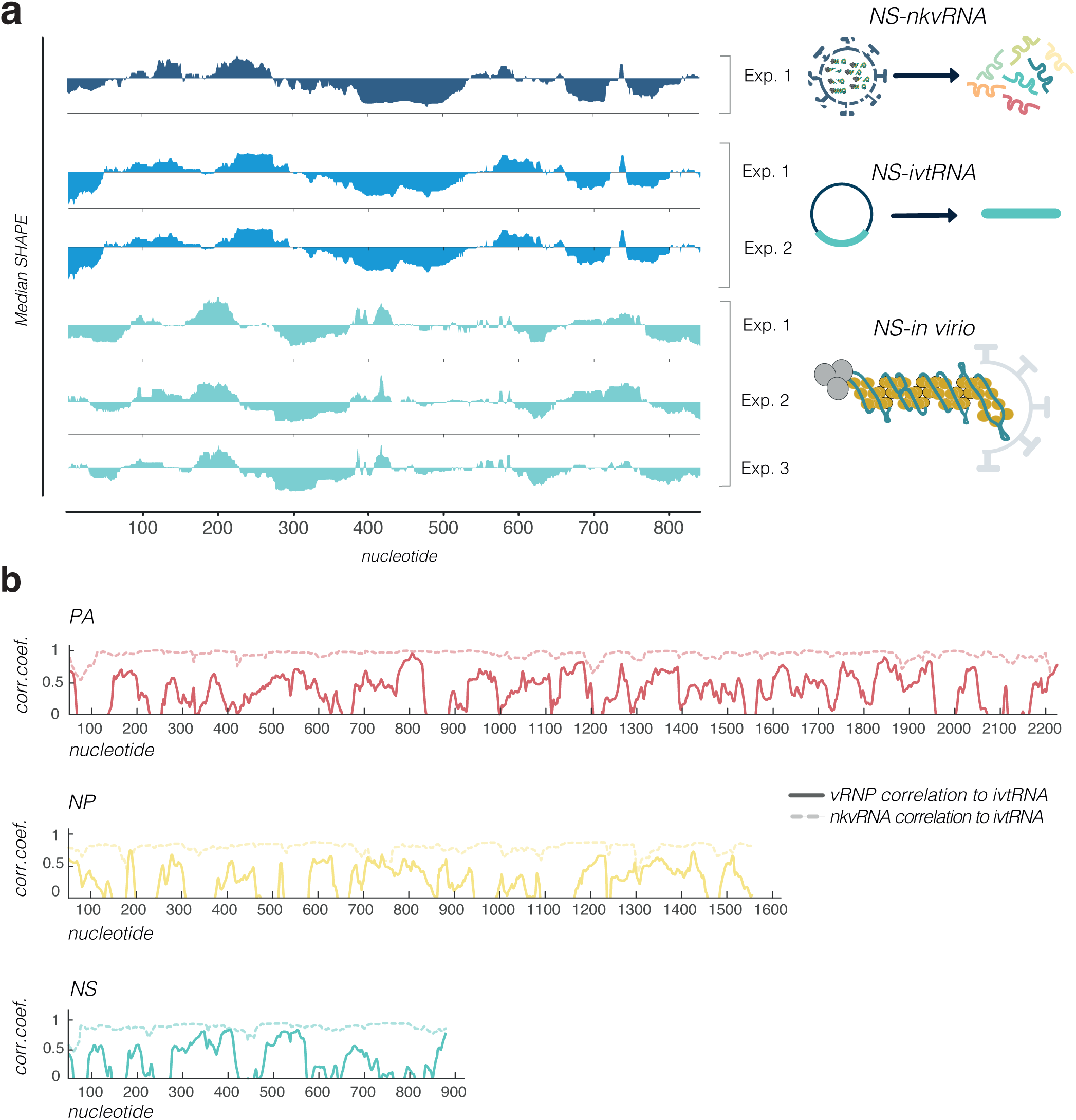
SHAPE reactivity variation between replicates and SHAPE reactivity correlation between different vRNA samples. **a,** Comparison of the median SHAPE reactivities of NS segment vRNA. Medians were calculated over 50 nucleotide windows and plotted relative to the global median. **b,** Sliding window correlation between *ex virio and in virio* vRNA samples. Pearson correlation was calculated over 50 nucleotide windows.

**Extended Data Figure 3.**
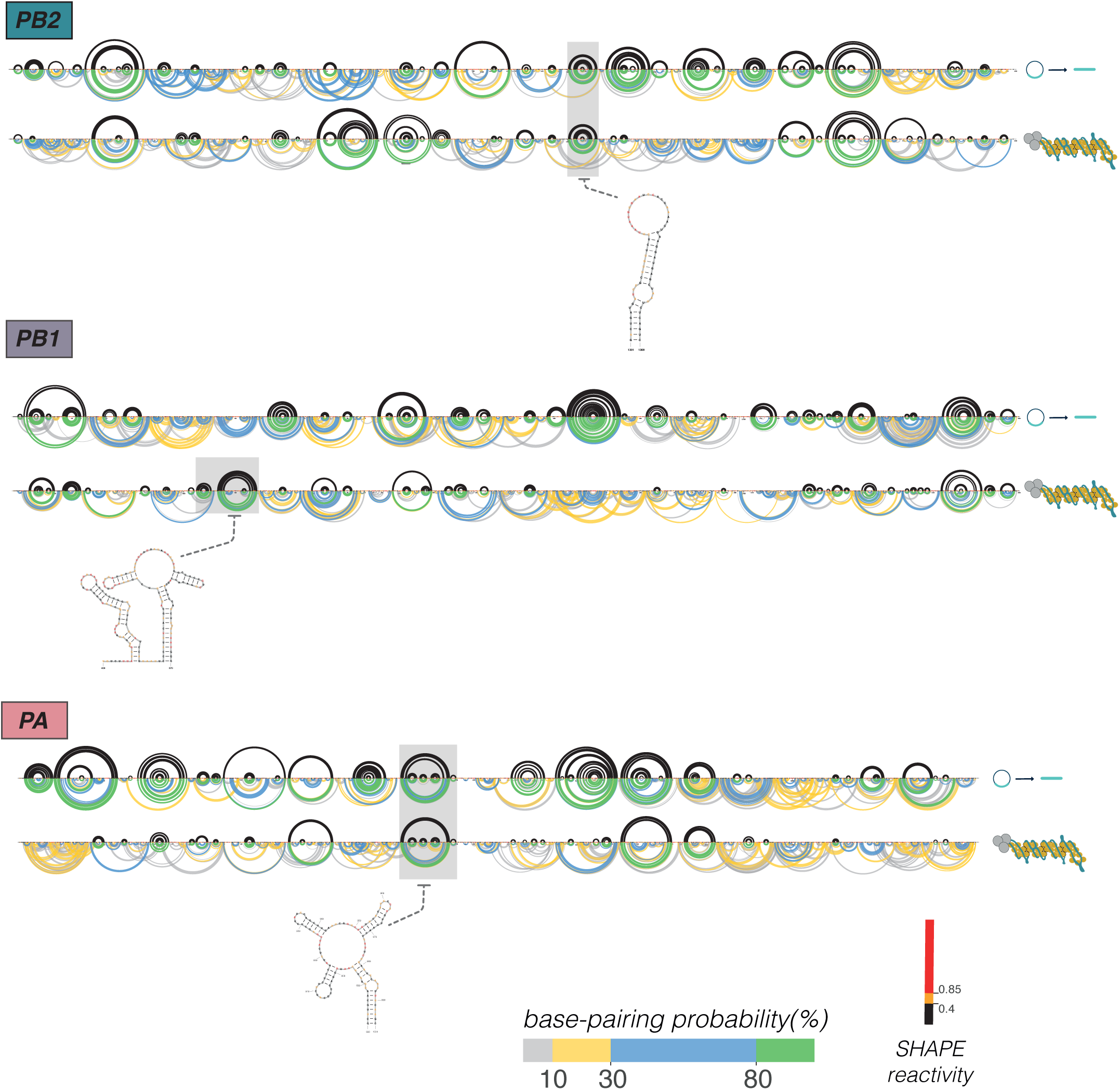
SHAPE-informed secondary RNA structure of the IAV polymerase segments. Black arcs indicate the maximum expected accuracy RNA structures; only the arcs associated with greater than 80% base-pairing probabilities are shown. Coloured arcs show base-pairing probabilities. Secondary structure examples of highlighted regions are shown.

**Extended Data Figure 4.**
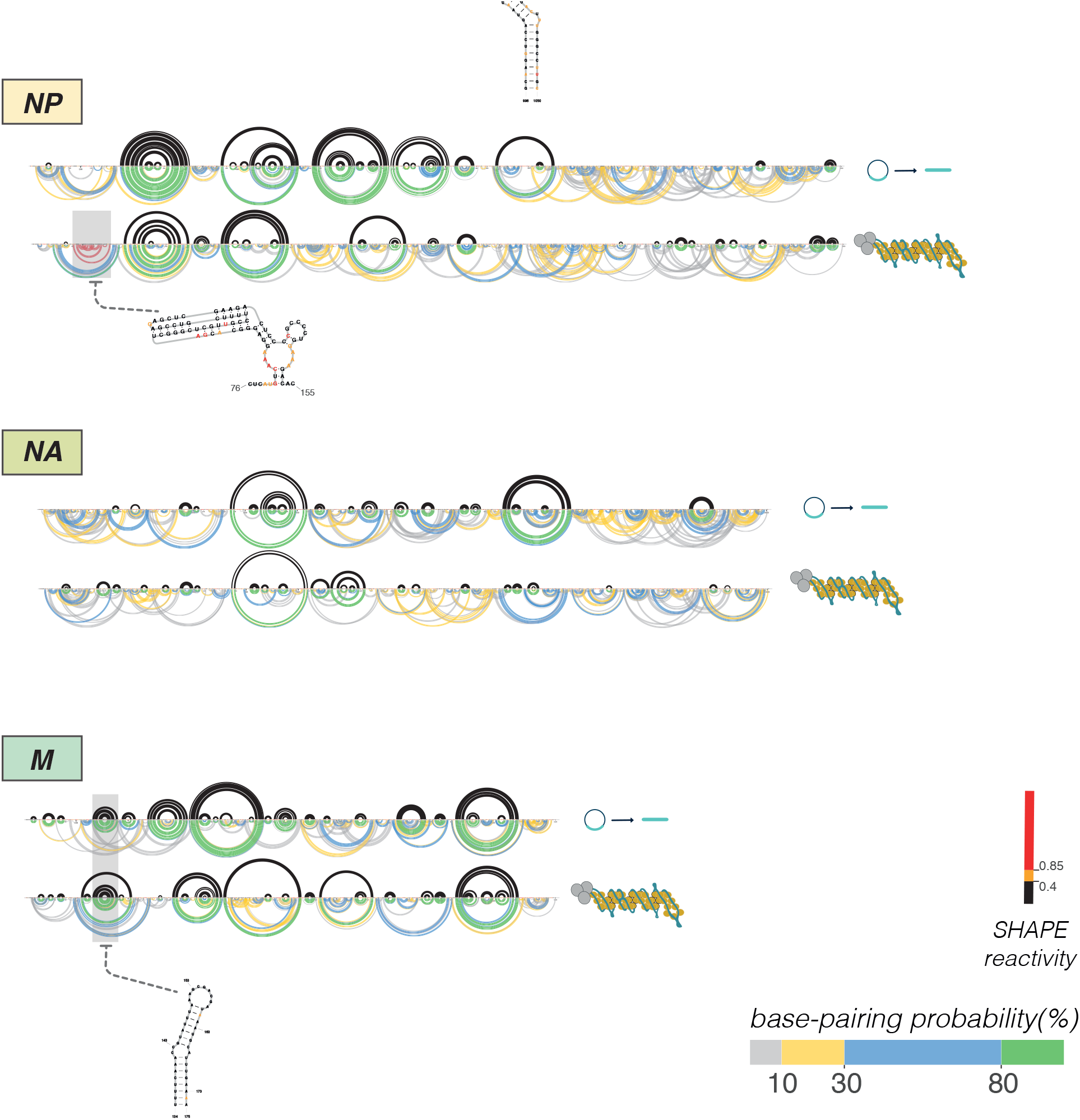
SHAPE-informed secondary RNA structure of the IAV HA, NP, NA and M segments. Black arcs indicate the maximum expected accuracy RNA structures; only the arcs associated with greater than 80% base-pairing probabilities are shown. Coloured arcs show base-pairing probabilities. Secondary structure examples of highlighted regions are shown. The pseudoknot in NP is highlighted in red.

**Extended Data Figure 5.**
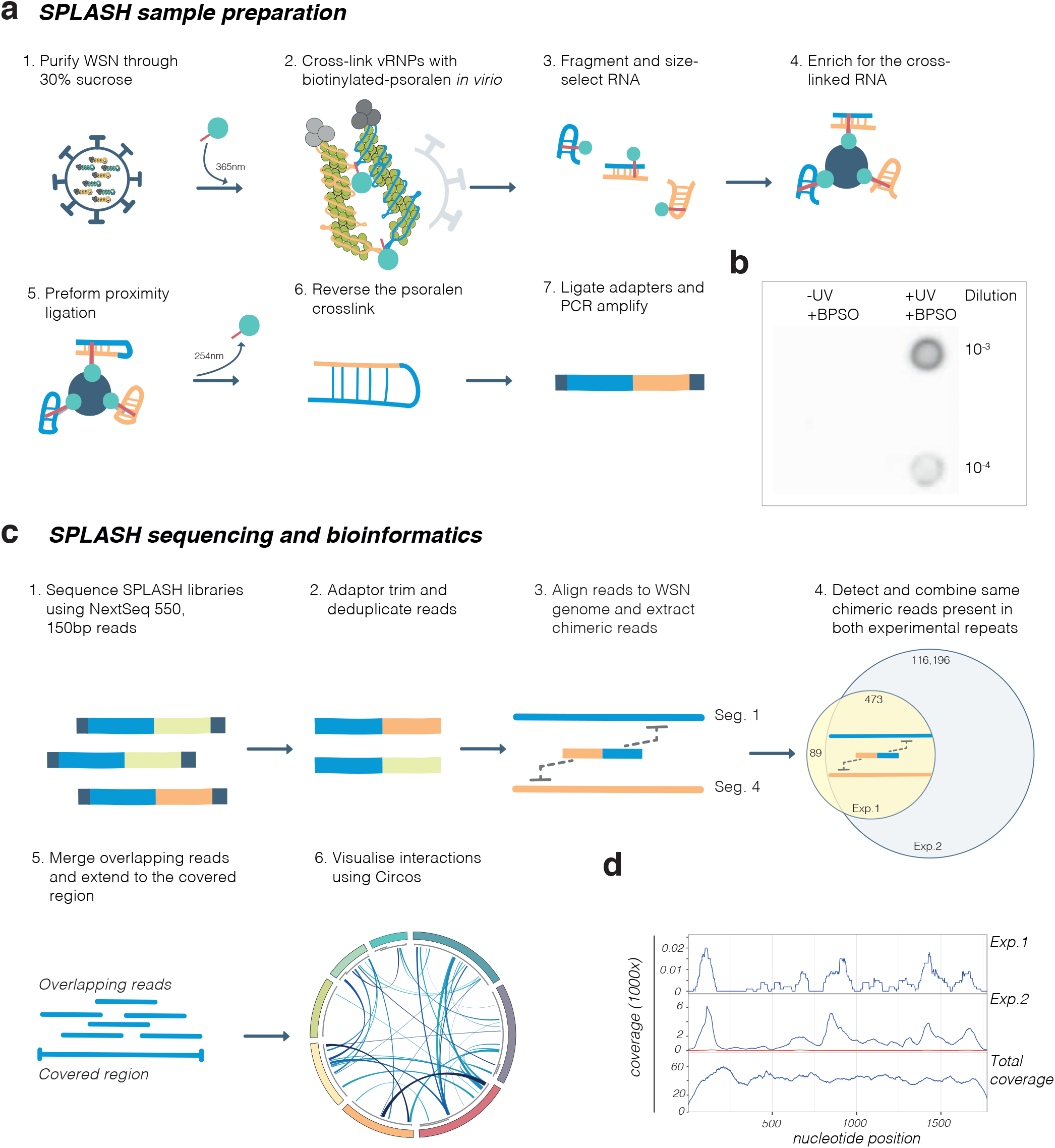
SPLASH samples and sequencing. **a,** Schematic showing SPLASH sample preparation. **b,** Anti-biotin dot blot analysis of RNA, crosslinked with biotinylated psoralen, extracted from viral particles. **c,** Schematic showing the SPLASH sequencing method and bioinformatics analysis steps. **d,** Chimeric reads aligned to the HA segment from two experimental replicates versus total RNA input coverage. Red trace indicates sample in which T4 RNA ligase I was omitted during proximity ligation (step 5 in **a**). BPSO, biotinylated psoralen; Exp., experiment.

**Extended Data Figure 6.**
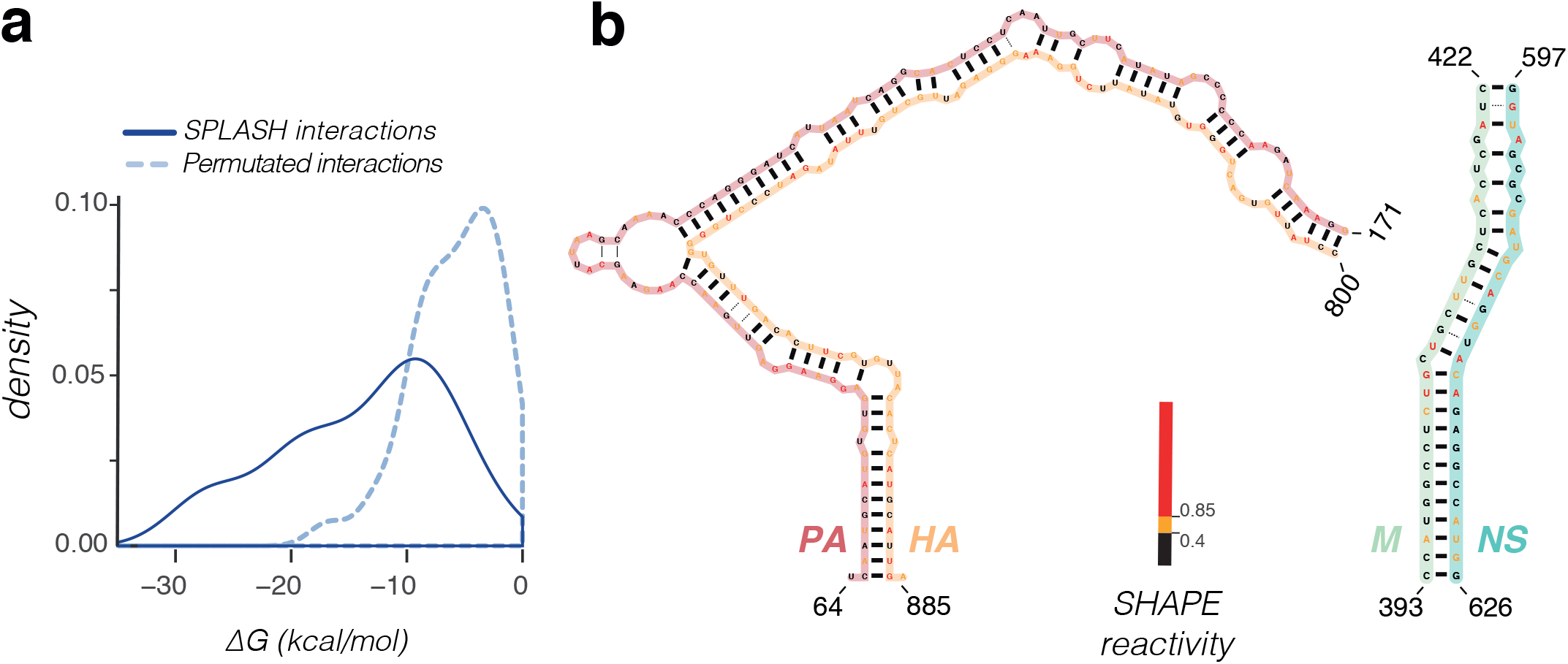
SHAPE-informed intermolecular RNA interactions. **a,** Probability distributions of the ΔG energies associated with the most common interactions identified by SPLASH versus a permutated interaction dataset as determined using SHAPE-informed structure prediction. **b,** Examples of intermolecular secondary RNA interaction structures based on SHAPE-informed structure predictions.

**Extended Data Figure 7.**
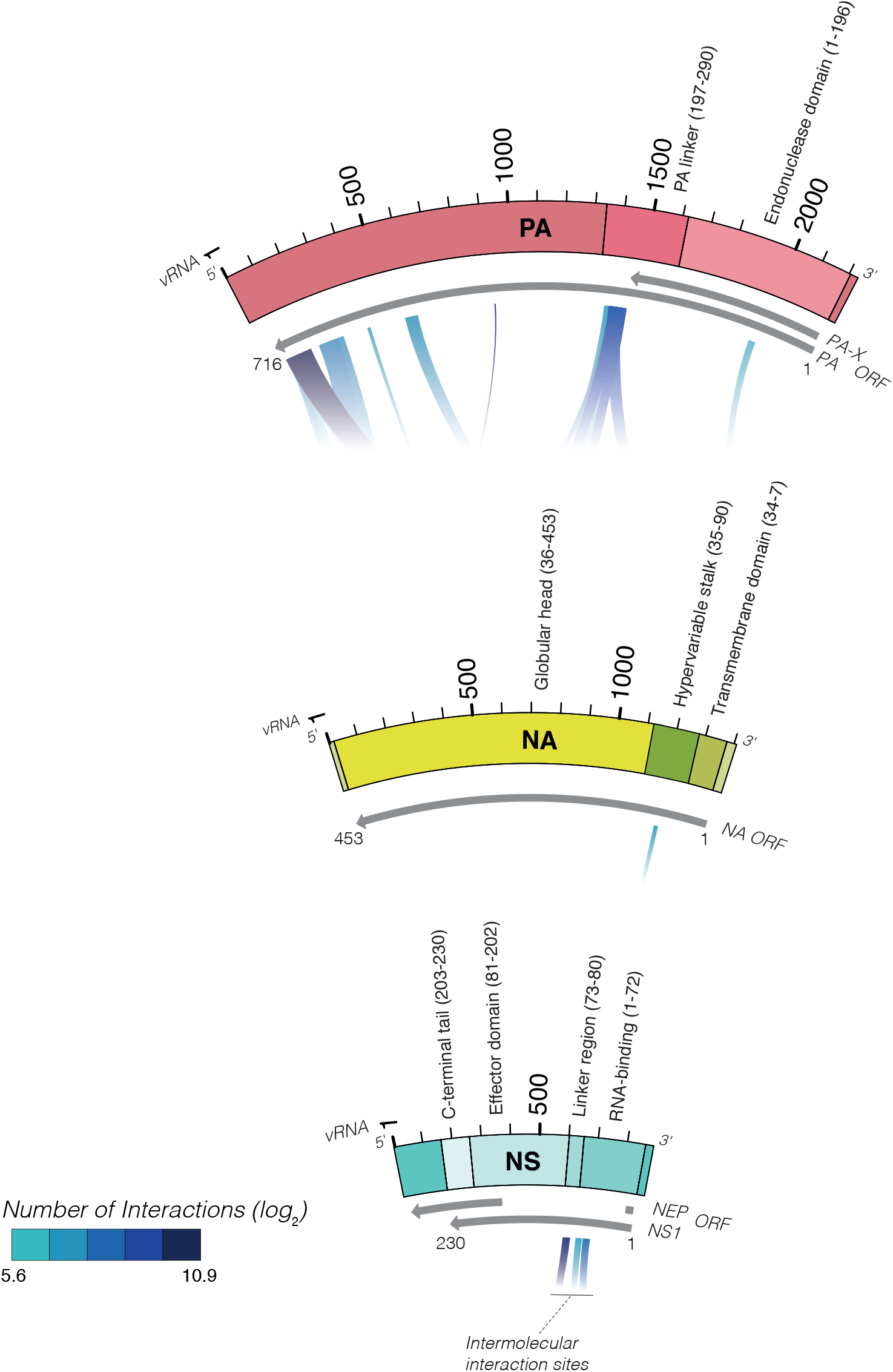
Intermolecular interactions associated with evolutionary flexible protein regions. Schematics of polymerase acidic protein, nonstructural protein 1 and neuraminidase protein. Numbers in the brackets indicate protein region amino acid positions. PA, polymerase acidic protein; NS1, nonstructural protein 1; NA, neuraminidase; ORF, open reading frame.

**Extended data table 1.**
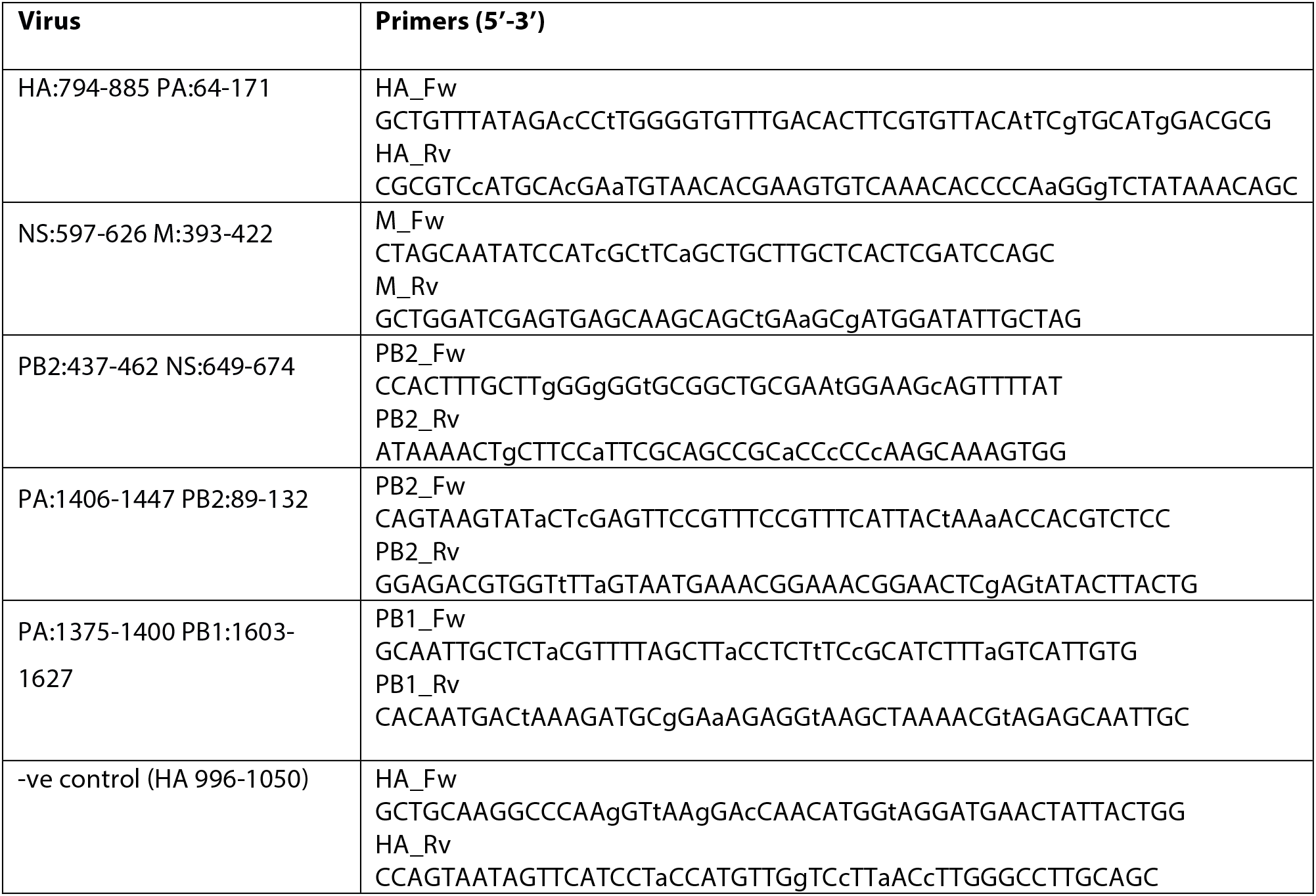
Primers used for synonymous mutagenesis of influenza segments. Mutated nucleotides are shown in lower case letters.

**Extended data table 2.**
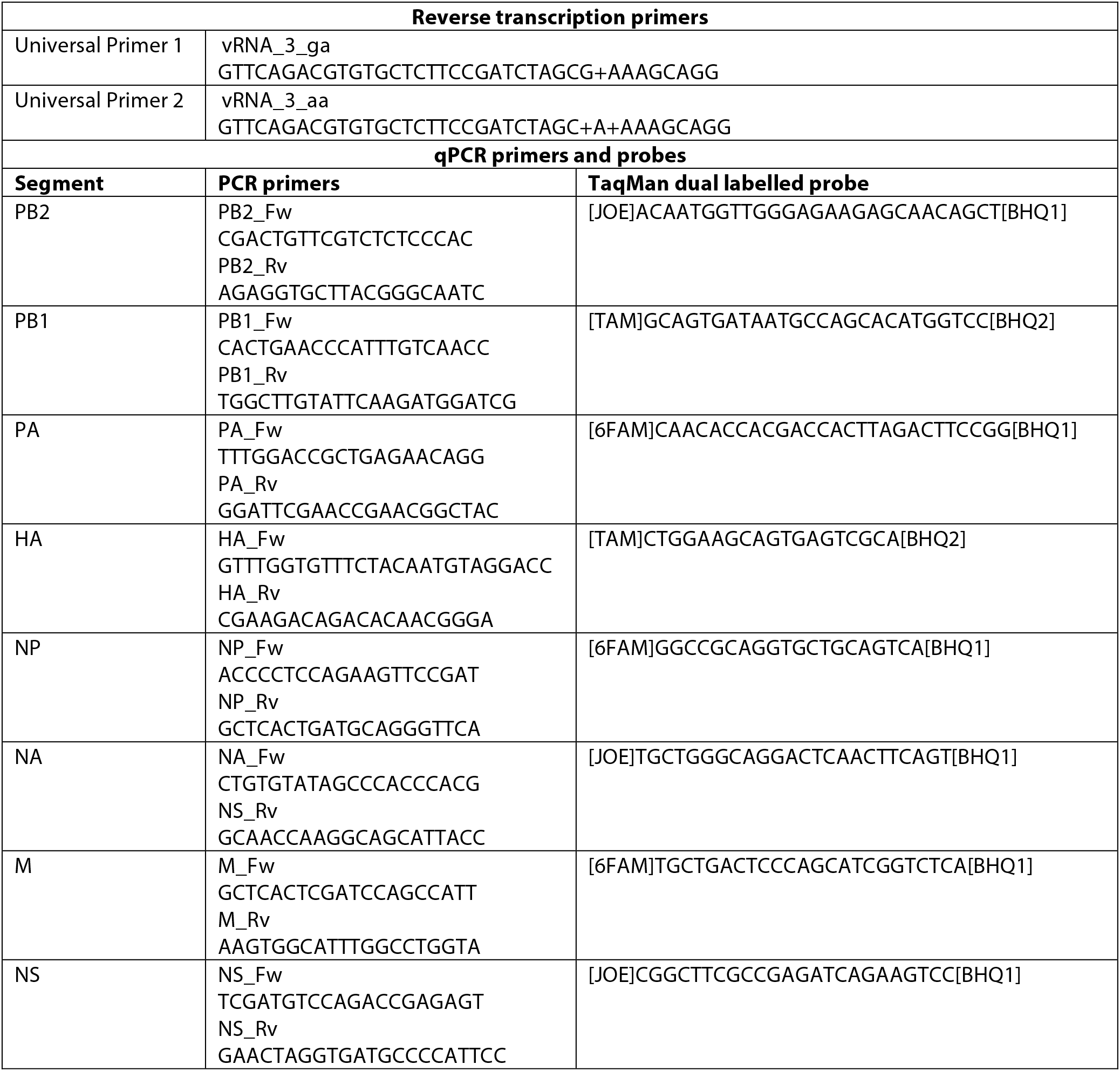
Primers and probes used for RT-qPCR. Locked nucleic acid nucleotides are indicated by + sign.

## References

1. Eisfeld AJ, Neumann G, Kawaoka Y. At the centre: Influenza A virus ribonucleoproteins. Nature Reviews Microbiology. 2015;13(1):28–41.

2. te Velthuis AJ, Fodor E. Influenza virus RNA polymerase: Insights into the mechanisms of viral RNA synthesis. Nature Reviews Microbiology. 2016;14(8):479–493.

3. Gerber M, Isel C, Moules V, Marquet R. Selective packaging of the influenza A genome and consequences for genetic reassortment. Trends Microbiol. 2014;22(8):446–455.

4. Hutchinson EC, von Kirchbach JC, Gog JR, Digard P. Genome packaging in influenza A virus. J Gen Virol. 2010;91 (2):313–328.

5. Lowen AC. Constraints, drivers, and implications of influenza A virus reassortment. Annual Review of Virology. 2017;4(1).

6. Taubenberger JK, Kash JC. Influenza virus evolution, host adaptation, and pandemic formation. Cell host & microbe. 2010;7(6):440–451.

7. Pflug A, Lukarska M, Resa-Infante P, Reich S, Cusack S. Structural insights into RNA synthesis by the influenza virus transcription-replication machine. Virus Res. 2017.

8. Arranz R, Coloma R, Chichon FJ, et al. The structure of native influenza virion ribonucleoproteins. Science. 2012;338(6114):1634–1637. doi: 10.1126/science.1228172 [doi].

9. Moeller A, Kirchdoerfer RN, Potter CS, Carragher B, Wilson IA. Organization of the influenza virus replication machinery. Science. 2012;338(6114):1631–1634. doi: 10.1126/science.1227270 [doi].

10. Wilkinson KA, Gorelick RJ, Vasa SM, et al. High-throughput SHAPE analysis reveals structures in HIV-1 genomic RNA strongly conserved across distinct biological states. PLoS biology. 2008;6(4):e96.

11. Smola MJ, Rice GM, Busan S, Siegfried NA, Weeks KM. Selective 2 [prime]-hydroxyl acylation analyzed by primer extension and mutational profiling (SHAPE-MaP) for direct, versatile and accurate RNA structure analysis. Nature protocols. 2015;10(11):1643–1669.

12. Baudin F, Bach C, Cusack S, Ruigrok RW. Structure of influenza virus RNP. I. influenza virus nucleoprotein melts secondary structure in panhandle RNA and exposes the bases to the solvent. EMBO J. 1994;13(13):3158–3165.

13. Kobayashi Y, Dadonaite B, van Doremalen N, Suzuki Y, Barclay WS, Pybus OG. Computational and molecular analysis of conserved influenza A virus RNA secondary structures involved in infectious virion production. RNA biology. 2016;13(9):883–894.

14. Gultyaev AP, Tsyganov-Bodounov A, Spronken MI, Van Der Kooij S, Fouchier RA, Olsthoorn RC. RNA structural constraints in the evolution of the influenza A virus genome NP segment. RNA biology. 2014;11 (7):942–952.

15. Lee N, Le Sage V, Nanni AV, Snyder DJ, Cooper VS, Lakdawala SS. Genome-wide analysis of influenza viral RNA and nucleoprotein association. Nucleic Acids Res. 2017;45(15):8968–8977.

16. Gavazzi C, Yver M, Isel C, et al. A functional sequence-specific interaction between influenza A virus genomic RNA segments. Proc Natl Acad Sci U S A. 2013;110(41):16604–16609. doi: 10.1073/pnas.1314419110 [doi].

17. Fournier E, Moules V, Essere B, et al. A supramolecular assembly formed by influenza A virus genomic RNA segments. Nucleic Acids Res. 2011;40(5):2197–2209.

18. Marsh GA, Rabadan R, Levine AJ, Palese P. Highly conserved regions of influenza a virus polymerase gene segments are critical for efficient viral RNA packaging. J Virol. 2008;82(5):2295–2304. doi: JVI.02267-07 [pii].

19. Gog JR, Afonso EDS, Dalton RM, et al. Codon conservation in the influenza A virus genome defines RNA packaging signals. Nucleic Acids Res. 2007;35(6):1897–1907.

20. Hutchinson EC, Wise HM, Kudryavtseva K, Curran MD, Digard P. Characterisation of influenza A viruses with mutations in segment 5 packaging signals. Vaccine. 2009;27(45):6270–6275.

21. Hutchinson EC, Curran MD, Read EK, Gog JR, Digard P. Mutational analysis of cis-acting RNA signals in segment 7 of influenza A virus. J Virol. 2008;82(23):11869–11879. doi: 10.1128/JVI.01634-08 [doi].

22. Aw JGA, Shen Y, Wilm A, et al. In vivo mapping of eukaryotic RNA interactomes reveals principles of higher-order organization and regulation. Mol Cell. 2016;62(4):603–617.

23. Greenbaum BD, Li OT, Poon LL, Levine AJ, Rabadan R. Viral reassortment as an information exchange between viral segments. Proc Natl Acad Sci U S A. 2012;109(9):3341–3346. doi: 10.1073/pnas.1113300109 [doi].

24. White MC, Steel J, Lowen AC. Heterologous packaging signals on segment 4, but not segment 6 or segment 8, limit influenza A virus reassortment. J Virol. 2017;91 (11):10.1128/JVI.00195-17. Print 2017 Jun 1. doi: e00195-17 [pii].

25. Gao Q, Palese P. Rewiring the RNAs of influenza virus to prevent reassortment. Proc Natl Acad Sci U S A. 2009;106(37):15891–15896. doi: 10.1073/pnas.0908897106 [doi].

26. Noda T, Sugita Y, Aoyama K, et al. Three-dimensional analysis of ribonucleoprotein complexes in influenza A virus. Nature communications. 2012;3:639.

27. Noda T, Sagara H, Yen A, et al. Architecture of ribonucleoprotein complexes in influenza A virus particles. Nature. 2006;439(7075):490–492.

28. Fournier E, Moules V, Essere B, et al. Interaction network linking the human H3N2 influenza A virus genomic RNA segments. Vaccine. 2012;30(51):7359–7367.

29. Fodor E, Devenish L, Engelhardt OG, Palese P, Brownlee GG, Garcia-Sastre A. Rescue of influenza A virus from recombinant DNA. J Virol. 1999;73(11):9679–9682.

30. Turner R, Shefer K, Ares M, Jr. Safer one-pot synthesis of the ‘SHAPE’ reagent 1-methyl-7-nitroisatoic anhydride (1m7). RNA. 2013;19(12):1857–1863. doi: 10.1261/rna.042374.113 [doi].

31. Wilkinson KA, Merino EJ, Weeks KM. Selective 2 [prime]-hydroxyl acylation analyzed by primer extension (SHAPE): Quantitative RNA structure analysis at single nucleotide resolution. Nature protocols. 2006;1 (3):1610–1616.

32. Aw JGA, Shen Y, Nagarajan N, Wan Y. Mapping RNA-RNA interactions globally using biotinylated psoralen. JoVE (Journal of Visualized Experiments). 2017(123):e55255–e55255.

33. Jiang H, Lei R, Ding S, Zhu S. Skewer: A fast and accurate adapter trimmer for next-generation sequencing paired-end reads. BMC Bioinformatics. 2014;15(1):182.

34. Martin M. Cutadapt removes adapter sequences from high-throughput sequencing reads. EMBnet.journal. 2011;17(1):pp. 10–12.

35. Dobin A, Davis CA, Schlesinger F, et al. STAR: Ultrafast universal RNA-seq aligner. Bioinformatics. 2013;29(1):15–21.

36. Krzywinski M, Schein J, Birol I, et al. Circos: An information aesthetic for comparative genomics. Genome Res. 2009;19(9):1639–1645. doi: 10.1101/gr.092759.109 [doi].

37. Busch A, Richter AS, Backofen R. IntaRNA: Efficient prediction of bacterial sRNA targets incorporating target site accessibility and seed regions. Bioinformatics. 2008;24(24):2849–2856.

38. Lorenz R, Bernhart SH, Zu Siederdissen CH, et al. ViennaRNA package 2.0. Algorithms for Molecular Biology. 2011;6(1):26.

39. Reuter JS, Mathews DH. RNAstructure: Software for RNA secondary structure prediction and analysis. BMC Bioinformatics. 2010;11(1):1.

